## supplemental_data for "Systems analysis of miR-199a/b-5p and multiple miR-199a/b-5p targets during chondrogenesis"

**Supplementary Material**

| log2FoldChange | Padj | name |
|---|---|---|
| 2.616518 | 0 | TGFBI |
| 2.709744 | 0 | TNC |
| 8.394896 | 0 | IGFBP5 |
| 3.538533 | 0 | DCN |
| 1.611538 | 0 | PGK1 |
| 5.15687 | 0 | ANGPTL4 |
| 2.829109 | 0 | IL1R1 |
| 5.613852 | 0 | TSC22D3 |
| -1.88916 | 0 | TUBA1B |
| 3.388879 | 0 | PLIN2 |
| -2.54946 | 0 | NRP1 |
| 2.354566 | 0 | P4HA2 |
| 2.77377 | 0 | COL7A1 |
| 2.607228 | 0 | UGP2 |
| 2.625552 | 0 | DUSP1 |
| 6.370805 | 0 | FKBP5 |
| 3.795323 | 0 | PPP1R3C |
| 2.315841 | 0 | BNIP3 |
| 3.197938 | 0 | SNAI2 |
| 5.970173 | 0 | MT1X |
| 2.234107 | 0 | SLC2A1 |
| 2.184884 | 0 | STEAP3 |
| -2.30296 | 0 | SHISAL1 |
| 6.928209 | 0 | COMP |
| 5.666913 | 0 | HILPDA |
| 3.367955 | 0 | APOL2 |
| 3.824101 | 0 | CRYAB |
| 5.081568 | 0 | PRG4 |
| 3.115528 | 0 | MXI1 |
| 2.436917 | 0 | FOXO3 |
| -2.85381 | 0 | COLEC12 |
| 5.293173 | 0 | BMP2 |
| -3.16176 | 0 | EXT1 |
| 3.374564 | 0 | SOX9 |
| 4.771362 | 0 | ARHGEF19 |
| 2.938684 | 0 | FAM162A |
| -3.75609 | 0 | ADAMTS5 |
| 4.84389 | 0 | FOXO1 |
| -4.97391 | 0 | CLDN11 |
| -3.92651 | 0 | CXCL12 |
| -5.09578 | 7.06E-304 | RGS4 |
| 2.825376 | 9.64E-298 | NMB |
| 2.813253 | 5.26E-297 | SLC6A8 |

| | | |
|---|---|---|
| -2.37377 | 2.83E-292 | SH2B3 |
| -1.99395 | 1.87E-288 | HAS2 |
| 2.708725 | 2.49E-287 | PFKFB4 |
| 2.119636 | 1.30E-283 | BGN |
| -1.83236 | 4.99E-277 | ANXA1 |
| 3.210302 | 1.70E-276 | SNAI1 |
| 2.847524 | 4.07E-275 | SAT1 |
| 3.967469 | 5.75E-275 | DEPP1 |
| 4.137285 | 1.10E-274 | SEC14L2 |
| 1.384284 | 4.58E-269 | SH3PXD2A |
| -2.5886 | 3.37E-267 | CDC42EP3 |
| 4.743967 | 1.10E-264 | ATP1B1 |
| -3.81618 | 3.82E-263 | DKK1 |
| -1.97544 | 2.30E-262 | BCL2L1 |
| 1.427985 | 3.96E-262 | ATP1A1 |
| -1.62554 | 1.10E-261 | COTL1 |
| 2.248828 | 2.02E-259 | PYGL |
| 1.666937 | 2.93E-258 | SPARC |
| 6.06187 | 7.90E-258 | ROS1 |
| 2.285569 | 9.94E-258 | KLF9 |
| 3.22922 | 1.16E-245 | TXNIP |
| 3.340382 | 1.65E-242 | DDIT4 |
| -2.96542 | 2.15E-242 | DCBLD2 |
| 1.948309 | 7.83E-242 | COL5A2 |
| 3.433647 | 6.86E-237 | GLUL |
| -2.75208 | 1.75E-235 | FHL2 |
| 2.079477 | 6.83E-235 | COL3A1 |
| 9.995813 | 1.32E-232 | SERPINA3 |
| 2.153327 | 9.18E-227 | TUT7 |
| 3.781249 | 1.48E-226 | COL11A1 |
| 2.563916 | 8.36E-224 | RHOB |
| -1.62741 | 3.09E-219 | CAPN2 |
| 3.888793 | 5.01E-219 | NFIL3 |
| 2.433366 | 3.06E-214 | ZNF395 |
| 2.802341 | 2.29E-213 | SAP30 |
| -2.12046 | 2.45E-212 | NQO1 |
| 3.171023 | 2.98E-205 | PLPP1 |
| 4.079239 | 7.13E-204 | MGP |
| 4.916167 | 4.66E-201 | MMP3 |
| 1.484742 | 2.22E-200 | ATF4 |
| 3.424358 | 3.38E-200 | ACSL1 |
| -1.94314 | 1.12E-198 | GLIPR1 |
| -1.45435 | 5.81E-197 | CFL1 |
| 6.270373 | 3.23E-194 | MAOA |
| 3.667951 | 3.31E-192 | TYRO3 |
| -2.55527 | 1.41E-190 | ADGRL4 |
| -2.3719 | 5.15E-189 | SLC20A1 |
| -2.53549 | 4.76E-188 | SCARA3 |

| | | |
|---|---|---|
| 2.350622 | 3.37E-186 | TIPARP |
| 2.674425 | 9.07E-186 | BTG1 |
| 2.042468 | 4.15E-183 | MIOS |
| 3.66087 | 9.66E-182 | NDUFA4L2 |
| 1.49663 | 1.46E-181 | FAM168A |
| -1.11214 | 1.57E-181 | CAV1 |
| -2.5923 | 4.55E-180 | CRIM1 |
| 1.043972 | 8.22E-180 | COL6A3 |
| 3.216222 | 3.72E-178 | PNRC1 |

**Supplementary Table 1.** Log 2 Fold Changes, BH adjusted P values and gene symbols from the first 100 most significantly (smallest adjusted P values) from performing differential expression analysis to contrast control/ undifferentiated chondrogenesis samples measured at day 1 of chondrogenesis against control/ undifferentiated chondrogenesis samples measured at day 0 of chondrogenesis.

| Gene | Barter D1/D0 | Barter D3/D0 | Barter D6/D0 | Barter D10/D0 | Barter D14/D0 |
|---|---|---|---|---|---|
| *ABHD17C* | NA | NA | NA | NA | NA |
| *ATP13A2* | -0.05 | -0.171 | -0.211 | -0.090 | **-0.341** |
| *CAV1* | **-3.380** | **-3.319** | **-1.522** | **-2.013** | **-1.998** |
| *CTSL* | NA | NA | NA | NA | NA |
| *DDR1* | **1.091** | **0.855** | **1.17** | **1.27** | **1.571** |
| *FZD6* | **-1.074** | **-0.781** | **-1.160** | **-1.087** | **-1.074** |
| *GIT1* | -0.007 | 0.071 | -0.156 | -0.118 | -0.044 |
| *HIF1A* | 0.075 | 0.646 | 0.280 | -0.193 | 0.075 |
| *HK2* | 0.141 | 0.401 | 0.111 | -0.051 | 0.141 |
| *HSPA5* | NA | NA | NA | NA | NA |
| *ITGA3* | **-2.289** | **-2.463** | **-2.063** | **-2.689** | **-2.289** |
| *M6PR* | **-0.687** | -0.264 | **-0.871** | -0.285 | **-0.697** |
| *MYH9* | **-1.756** | **-1.742** | **-2.554** | **-2.105** | **-1.756** |
| *NECTIN2* | NA | NA | NA | NA | NA |
| *NINL* | **-0.511** | **0.406** | **-0.364** | **-0.493** | **-0.511** |
| *PDE4D* | **-0.283** | **-0.406** | 0.082 | -0.152 | **-0.283** |
| *PLXND1* | **0.322** | **0.313** | **0.498** | **0.309** | **0.322** |
| *PXN* | -0.786 | -0.228 | -0.148 | -0.172 | -0.168 |
| *SLC35A3* | **0.996** | **0.791** | **1.314** | **1.212** | **0.996** |
| *SLC35D1* | **1.483** | **1.391** | **1.665** | **1.652** | **1.483** |
| *VPS26A* | NA | NA | NA | NA | NA |

**Supplementary Table 2.** Log2fc values from the chondrogenesis microarray study. The 21 genes shown were found through bioinformatic analysis of the RNAseq data. Values that are bold were significantly differentially expressed (adjusted P value is less than 0.05). Red highlighted values were down-regulated and blue highlighted genes were up-regulated. NA means the gene were not found in the study.

| Gene | Huynh D1/D0 | Huynh D3/D1 | Huynh D7/D3 | Huynh D14/ D7 | Huynh D21/ D7 | Huynh D21/ D14 | Huang MSC28/ MSC0 | Huang c28/ MSC0 | Huang c28/ MSC28 |
|---|---|---|---|---|---|---|---|---|---|
| *ABHD17C* | **-1.15** | -0.417 | 0.107 | -0.373 | -0.173 | 0.201 | -0.222 | -0.214 | 0.007 |
| *ATP13A2* | -0.387 | -0.698 | **0.577** | -0.06 | -0.208 | -0.14 | -0.07 | 0.355 | 0.425 |
| *CAV1* | **-0.99** | -0.221 | **-1** | -0.323 | 0.0429 | 0.366 | -0.572 | -0.207 | 0.365 |
| *CTSL* | **-0.957** | -0.0991 | -0.518 | -0.279 | -0.177 | 0.102 | NA | NA | NA |
| *DDR1* | 1.18 | -0.681 | -0.145 | 0.076 | 0.2 | 0.123 | 0.395 | **0.636** | 0.241 |
| *FZD6* | -0.362 | **-0.763** | **-0.367** | 0.104 | **0.41** | 0.306 | NA | NA | NA |
| *GIT1* | 0.188 | 0.0364 | **-0.352** | -0.354 | **-0.48** | -0.126 | **-0.472** | -0.354 | 0.238 |
| *HIF1A* | -0.686 | 0.011 | 0.059 | **0.581** | **0.759** | 0.178 | **0.56** | 0.344 | -0.216 |
| *HK2* | 0.556 | -0.212 | **-1.99** | **-0.645** | -0.552 | 0.093 | -0.325 | **0.832** | **1.16** |
| *HSPA5* | **2.89** | -0.443 | **-1.01** | **-1.42** | **-1.51** | -0.089 | NA | NA | NA |
| *ITGA3* | **-1.57** | **-1.01** | -0.26 | -0.729 | -0.019 | 0.709 | **-2.94** | **-4.78** | **-1.84** |
| *M6PR* | -0.269 | 0.056 | -0.005 | 0.0286 | -0.086 | -0.115 | **-0.775** | **-0.991** | -0.216 |
| *MYH9* | -0.236 | **-0.833** | 0.276 | -0.144 | **-0.427** | -0.283 | **-1.69** | **-0.952** | **0.734** |
| *NECTIN2* | NA | NA | NA | NA | NA | NA | NA | NA | NA |
| *NINL* | **-1.18** | **0.513** | 0.176 | -0.153 | 0.133 | -0.02 | NA | NA | NA |
| *PDE4D* | 0.909 | **-1.01** | 0.278 | -0.162 | 0.425 | 0.587 | **0.605** | **0.684** | 0.078 |
| *PLXND1* | **-0.608** | **-0.81** | 0.039 | 0.139 | 0.173 | 0.034 | **-1.46** | **-1.59** | -0.134 |
| *PXN* | **-1.01** | **-0.599** | 0.02 | -0.226 | -0.123 | 0.103 | NA | NA | NA |
| *SLC35A3* | 0.424 | -0.278 | -0.278 | 0.175 | 0.343 | 0.168 | NA | NA | NA |
| *SLC35D1* | 0.381 | -0.143 | 0.616 | 0.764 | **0.798** | 0.0345 | **-1.18** | 0.158 | **1.34** |
| *VPS26A* | -0.949 | -0.364 | -0.278 | -0.397 | -0.114 | 0.283 | -0.773 | -1.19 | -0.422 |

**Supplementary Table 3.** Log2fc values from the 21 *miR-199a/b-5p* targets from bioinformatic analysis of the RNAseq data. Results from *Huynh et al (2019)* and *Huang et al (2010)* are shown here as these are the datasets found in the *SkeletalVis* repository which have the keywords "Chondrogenesis" and "MSC differentiation" [1,2]. Significantly differentially expressed genes with an adjusted P value of less than 0.05 are bold and either red if they were down-regulated or blue if they were up-regulated. NA means the gene was not found in the study. Data from *Huynh et al (2019)* consisted of time points: days 0 1, 3, 7, 14 and 21 after chondrogenesis initiation and the data underwent step-wise DE analysis in *SkeletalVis*. Data from *Huang et al (2010)* consisted of chondrogenesis (c) or MSC samples measured at days 0 or 28, and these have been contrasted in *SkeletalVis*.

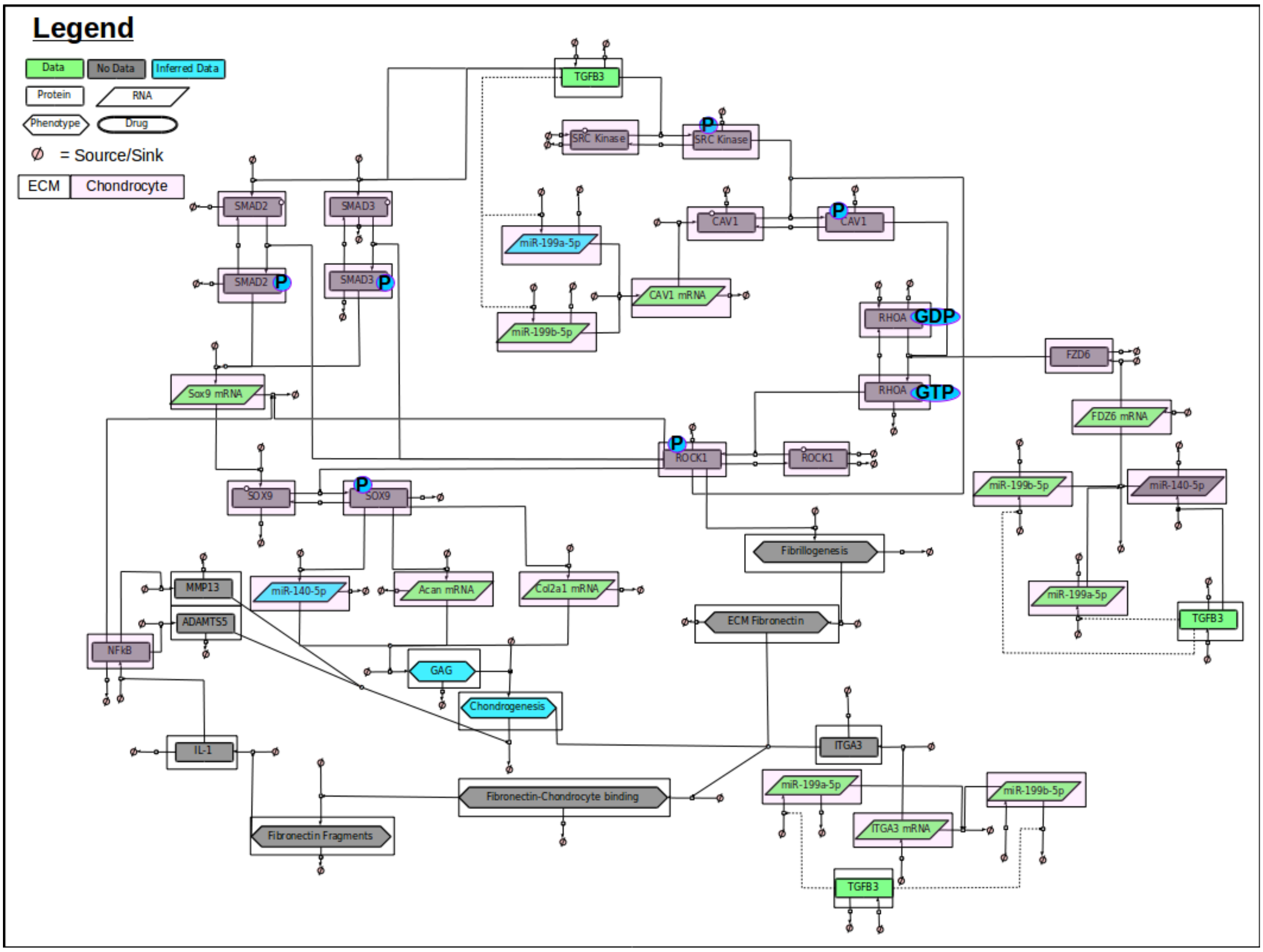

**Supplementary Figure 1.** GRN containing the broader scope of the biological system modelled in this paper. This GRN contains This broader GRN contained 40 species including: twelve proteins (TGFB3, SRC Kinase, CAV1, FZD6, ITGA3, SOX9, IL-1, NfkB, MMP13, ADAMTS5, SMAD2, SMAD3), six phospho-protein (phospho-SRC kinase, phospho-SOX9, phospho-CAV1, phospho-ROCK1, phospho-SMAD2, phospho-SMAD3), two RHOA states (GDP-RHOA, GTP-RHOA), six mRNAs (*SOX9*, *ACAN*, *COL2A1, CAV1, FZD6, ITGA3*), three miRNAs (*miR-140-5p*, *miR-199a-5p*, *miR-199b-5p*), and six phenotypes/ ECM constructs (GAG, Chondrogenesis, Fibrillogenesis, ECM fibronectin, Fibronectin-Chondrocyte binding, Fibronectin fragments). Each species as a sink and a source. Species are also shaped based on their properties: Proteins are rectangles, RNAs are rhombus, Phenotypes are hexagons and Drugs are oval. Species are also highlighted with a white box if they are found in the ECM or pink if they are found within a chondrocyte. Species are also colour coded: green if there is associated data, blue if there is some data and the rest has been inferred based on literature, or grey if there is no data associated with the species.

| Primer | Sequence |
|---|---|
| SOX9 F | ACTTGCACAACGCCGAG |
| SOX9 R | CTGGTACTTGTAATCCGGGTG |
| COL2A1 F | AACCAGATTGAGAGCATCCG |
| COL2A1 R | ACCTTCATGGCGTCCAAG |
| ACAN F | AGCGAGTTGTCATGGTCTG |
| ACAN R | TGTGGGACTGAAGTTCTTGG |
| FZD6 F | GAAGCAAAAAGACATGCACAGA |
| FZD6 R | TTCGACTTTCACTGATTGGATCT |
| ITGA3 F | GAGGACATGTGGCTTGGAGT |
| ITGA3 R | GTAGCGGTGGGCACAGAC |
| CAV1 F | ACAGCCCAGGGAAACCTC |
| CAV1 R | CGGATGGGAACGGTGTAG |
| hFZD6 IFC F | GCTCGCTAGCCTCGATCTCTCGTTACTCAGAAGCAAA |
| hFZD6 IFC R | CGACTCTAGACTCGATGGCACTAATATCGCTATCACAC |
| hITGA3 IFC F | GCTCGCTAGCCTCGACGGACCCGCTATTATCAGATC |
| hITGA3 IFC R | CGACTCTAGACTCGACTGGGAGCTGTTTATTGGTCG |
| hFZD6 mut F | GTGCATAGGTCACTTCGACTCTAACACAAATTTGCTTCTGAGTAACGAGA |
| hFZD6 mut R | TCTCGTTACTCAGAAGCAAATTTGTGTTAGAGTCGAAGTGACCTATGCAC |
| hITGA3 mut1 F | GTGGCTCAAGATGGATCGACTCTGAAAGGGGGAGGTGTC |
| hITGA3 mut1 R | GACACCTCCCCCTTTCAGAGTCGATCCATCTTGAGCCAC |

| Probe |
|---|
| 5'-FAM-TCTGGAGACTTCTGAACGAGAGCGA-IABkFQ-3' |
| 5'-FAM-AGACCTGAAACTCTGCCACCCTG-IABkFQ-3' |
| 5'-FAM-CTGGGTTTTCGTGACTCTGAGGGT-IABkFQ-3' |

**Supplementary Table 4.** Sequences of primers and probes used for mutagenesis experiments and knock-downs.

**Model Specifics for the initial chondrogenesis model**

**Initial conditions**

| Model species | Value (mmol/ml) | Description |
|---|---|---|
| *Acan* mRNA | 6.257096819 | Chondrogenesis biomarker. |
| *CAV1* mRNA | 13.21139434 | General genetic activity of CAV1 mRNA and protein. |
| *Col2a1* mRNA | 6.40998058 | Chondrogenesis biomarker. |
| *FZD6* mRNA | 8.692679393 | General genetic activity of FZD6 mRNA and protein. |
| GAG | 1 | Chondrogenesis biomarker. |
| HP199a | 0 | Drug reduces miR-199a-5p when event is triggered. |
| HP199b | 0 | Drug reduces miR-199b-5p when event is triggered. |
| *ITGA3* mRNA | 8.925880317 | General genetic activity of ITGA3 mRNA and protein. |
| miR-199a-5p | 10.5401385 | miRNA which modulates FZD6, ITGA3 and CAV1. |
| miR-199b-5p | 5.423464 | miRNA which modulates FZD6, ITGA3 and CAV1. |
| *SOX9* mRNA | 8.76039807 | Chondrogenesis biomarker. |

**Events**

In this kinetic model we have four events (HP199a activity, HP199a inactivity, HP199b activity, HP199b inactivity) have been used to simulate behaviours seen in our *miR-199a-5p* and *miR-199b-5p* knockdown experiments. When triggered, HP199a activity and HP199a inactivity both together lead to HP199a = 1 until time reaches 7 days, at which point HP199a = 0. Likewise, when both HP199b activity and HP199b inactivity are triggered, they lead to lead to HP199b = 1 until time reaches 7 days, at which point HP199a = 0. HP199a and HP199b will reduce their target miRNA by 90%-95% until day 7. HP199a and HP199b modulate global quantities which are fixed at 1. When not triggered the global quantities will not be modulated.

**Ordinary Differential Equations (ODEs)**

This was an ODE based kinetic model. Each of the model species have inputs and outputs which modulate their behaviours over the 14-day time course. All parameters have been rounded up to three decimal places. ch = chondrocyte compartment. ecm = extracellular matrix. ch and ecm were compartments, both equalled 1 so performed no modulation.

$$\frac{d[MIR199b\_5p].ch}{dt} = [+\,ch\,.\,4.60528$$

$$-\mathrm{ch}.\,(20\,.\,[MIR199b\_5p].\,[\mathrm{HP199b}])$$

$$-\,c\mathrm{h}.\,(0.505.\,[\mathrm{MIR199b\_5p}])\;]$$

$$\frac{d[MIR199a\_5p].ch}{dt} = [+\,ch\,.\,104.738$$

$$-\mathrm{ch}.\,(201.443\,.\,[MIR199a\_5p].\,[\mathrm{HP199a}])$$

$$-\,c\mathrm{h}.\,(9.066\,.\,[\mathrm{MIR199a\_5p}])\;]$$

$$\frac{d[ACAN\_mRNA].ch}{dt} = [+\,ch\,.\,(400.198\,.\,[SOX9\_mRNA])$$

$$-\mathrm{ch}.\,(507.529.\,[ACAN_{mRNA}]\;)]$$

$$\frac{d[COL2A1\_mRNA].ch}{dt} = [+\,ch\,.\,(778.139\,.\,[SOX9\_mRNA])$$

$$-\mathrm{ch}.\,(693.578.529.\,[COL2A1\_mRNA])\;]$$

$$\frac{d[SOX9\_mRNA].ch}{dt} = [+\,ch\,.\,5000$$

$$-\mathrm{ch}.\,(10\,.\,[SOX9\_mRNA]\,.\,[\mathrm{FZD6\_mRNA}]\;)$$

$$-\mathrm{ch}.\,([SOX9\_mRNA].\,[\mathrm{ITGA3\_mRNA}])$$

$$-\mathrm{ch}.\,([\mathrm{SOX9\_mRNA}]\,.\,[CAV1\_mRNA])]$$

$$\frac{d[FZD6\_mRNA].ch}{dt} = [+\,ch\,.\,(2051.29)$$

$$-\mathrm{ch}.\,(10.\,[FZD6\_mRNA]\,.\;\;[MIR199a\_5p])$$

$$-\,\mathrm{ch}.\,(18\,.\,[\mathrm{FZD6\_mRNA}]\,.\;\;[\mathrm{MIR199b\_5p}])]$$

$$\frac{d[ITGA3\_mRNA].ch}{dt} = [+\,ch\,.\,(1732.04)$$

$$-\mathrm{ch}.\,(10.\,[ITGA3\_mRNA]\,.\;\;[MIR199a\_5p])$$

$$-\,\mathrm{ch}.\,(18\,.\,[ITGA3\mathrm{\_mRNA}]\,.\;\;[\mathrm{MIR199b\_5p}])]$$

$$\frac{d[CAV1\_mRNA].ch}{dt} = [+ ch\,.\,(1732.04)$$
$$-\text{ch}.\,(10.\,[CAV1\_mRNA]\,.\ [MIR199a\_5p])$$
$$-\ \text{ch}.\,(18\,.\,[CAV1\_\text{mRNA}]\,.\ [\text{MIR199b\_5p}])]$$

**Model specifics for the enhanced chondrogenesis model**

**Initial conditions**

| Model species | Value (mmol/ml) | Description |
|---|---|---|
| *Acan* mRNA | 6.257096819 | Chondrogenesis biomarker. |
| CAV1 | 13.21139434 | General genetic activity of CAV1 mRNA and protein. |
| *Col2a1* mRNA | 6.40998058 | Chondrogenesis biomarker. |
| FZD6 | 8.692679393 | General genetic activity of FZD6 mRNA and protein. |
| GAG | 1 | Chondrogenesis biomarker. |
| HP199a | 0 | Drug reduces *miR-199a-5p* when event is triggered. |
| HP199b | 0 | Drug reduces *miR-199b-5p* when event is triggered. |
| ITGA3 | 8.925880317 | General genetic activity of ITGA3 mRNA and protein. |
| *miR-140-5p* | 6.5582925 | miRNA which modulates FZD6 and biomarker. |
| *miR-199a-5p* | 10.5401385 | miRNA which modulates FZD6, ITGA3 and CAV1. |
| *miR-199b-5p* | 5.423464 | miRNA which modulates FZD6, ITGA3 and CAV1. |
| OtherTargets | 25 | Representative of other *199a/b* targets. |
| OtherTargetsRegulators | 1000 | Regulator of OtherTargets to help modulation. |
| *SOX9* mRNA | 8.76039807 | Chondrogenesis biomarker. |
| SOX9PhosphoProtein | 0 | Promotes *ACAN*, *COL2A1*, *miR-140-5p.* |
| SOX9 | 1 | Promotes SOX9PhosphoProtein. |
| SRC | 1000 | Modulator of CAV1. |
| TGFB3 | 10000 | Represents chondrogenesis initiation. |

**Events**

In this kinetic model we have four events (HP199a activity, HP199a inactivity, HP199b activity, HP199b inactivity) have been used to simulate behaviours seen in our *miR-199a-5p* and *miR-199b-5p* knockdown experiments. When triggered, HP199a activity and HP199a inactivity both together lead to HP199a = 1 until time reaches 4.5 days, at which point HP199a = 0. Likewise, when both HP199b activity and HP199b inactivity are triggered, they lead to lead to HP199b = 1 until time reaches 4.5 days, at which point HP199a = 0. HP199a and HP199b will reduce their target miRNA by 90%-95% until day 4.5. HP199a and HP199b modulate global quantities which are fixed at 1. When not triggered the global quantities will not be modulated.

**Ordinary Differential Equations (ODEs)**

This was an ODE based kinetic model. Each of the model species have inputs and outputs which modulate their behaviours over the 14-day time course. All parameters have been rounded up to three decimal places. ch = chondrocyte compartment. ecm = extracellular matrix. ch and ecm were compartments, both equalled 1 so performed no modulation.

$$\frac{d[ACAN\ mRNA].ch}{dt} = [+\ ch.(\frac{100.[Sox9PhosphoProtein]}{1+\frac{1}{[Sox9PhosphoProtein]}}$$
$$-\ c\text{h}.(4.263.[\text{ACAN}\ mRNA])\ ]$$

$$\frac{d[CAV1].ch}{dt} = [+ch.\frac{152.229\ .9625.57.[\text{SRC}]}{3.(0.416+[SRC])+9625.57\ .(0.1+[SRC])}$$
$$-\ ch.\frac{12.892.[\text{CAV 1}]}{0.0972+[\text{CAV 1}]+0.0971\ .\frac{[miR-199a-5p]}{0.1318}}\ .[miR-199a-5p]$$
$$-\ ch.\frac{18.746.[\text{CAV 1}]}{0.0996+[\text{CAV 1}]+0.0996\ .\frac{[miR-199b-5p]}{0.057}}\ .[miR-199b-5p]$$
$$-\ c\text{h}.(0.267.[\text{CAV1}])\ ]$$

$$\frac{d[COL2A1\ mRNA].ch}{dt} = [+ch.(\frac{94.624\ .\ [Sox9PhosphoProtein]}{1+\frac{1}{[Sox9PhosphoProtein]}})$$

$$-ch\ .(3.005\ .[COL2A1\ mRNA]]$$

$$\frac{d[GAG].ecm}{dt} = [+\mathrm{ch}.(\frac{0.179.[\mathrm{COL2A1\ mRNA}]}{1+89.324}/[\mathrm{COL2A1\ mRNA}])$$

$$+ch.(\frac{4.70977.[\mathrm{ACAN\ mRNA}]}{1+1}/[\mathrm{ACAN\ mRNA}])$$

$$+ch/(\frac{3.97.[\mathrm{miR}-140-5\mathrm{p}]}{1+5}/[\mathrm{miR}-140-5\mathrm{pmRNA}])]$$

$$\frac{d[miR-140-5p].ch}{dt} = [+ch.(6.085.[\mathrm{Sox9PhosphoPrortein}])$$

$$-ch.(0.376.[\mathrm{miR}-140-5\mathrm{p}])]$$

$$\frac{d[miR-199a-5p].ch}{dt} = [+ch.\frac{100.41.\frac{[TGFB3]}{104.984}}{1+\frac{[TGFB3]}{104.984}/]}$$

$$-ch.(8.678.[\mathrm{miR}-199\mathrm{a}-5\mathrm{p}])$$

$$-ch.\left(\frac{(1.009.[\mathrm{miR}-199\mathrm{a}-5\mathrm{p}]}{0.017}\right).[HP199a]]$$

$$\frac{d[miR-199b-5p].ch}{dt} = [+ch.\frac{8.013\ .\frac{[TGFB3]}{0.015}}{1+\frac{[TGFB3]}{0.015}/]}$$

$$-ch.(0.90175.[\mathrm{miR}-199\mathrm{b}-5\mathrm{p}])$$

$$-ch.\left(\frac{(1.004.[\mathrm{miR}-199\mathrm{b}-5\mathrm{p}]}{0.124}\right).[HP199b]]$$

$$(d[OtherTargets].ch)/dt$$

$$= [+ch.\frac{1354.23\ .1554.29.[\mathrm{OtherTargetsRegulator}]}{0.008\ .(0.197+[\mathrm{OtherTargetsRegulator}])+1553.9\ .(0.009+[\mathrm{OtherTargetsRegulator}])})$$

$$-ch\ .\left(\frac{100.728\ .[\mathrm{OtherTargets}]}{0.1+[\mathrm{OtherTargets}]+0.1\frac{[\mathrm{miR}-199\mathrm{a}-5\mathrm{p}]}{0.103}}\right)$$

$$-ch\,.\,(\frac{121.391\ .\ [\mathrm{OtherTargets}]}{0.095+[\mathrm{OtherTargets}]+0.095\,\frac{[\mathrm{miR}-199\mathrm{b}-5\mathrm{p}]}{0.105}})$$
$$-ch.\,(0.097.\,[OtherTargets])]$$

$$\frac{d[OtherTargetsRegulator].\,ch}{dt}=[+ch\,.\,0.099$$
$$-ch.\,(8.652.\,[\mathrm{OtherT\ argetsRegulators}])]$$

$$\frac{d[SRC].\,ch}{dt}=[\ +ch.\,(\frac{[TGFB].\,0.117}{100})$$
$$-ch.\,1.407\ .\,[SRC]]$$

$$\frac{d[SOX9\ mRNA]}{dt}=[+ch.\,(604.499\ .\,[SOX9\ mRNA])$$
$$+ch.\,(2.1577.\,[\mathrm{CAV1}])$$
$$+ch.\,(1.551.\,[\mathrm{TGFB3}])$$
$$+ch.\,(8.44.\,[\mathrm{miR}-140-5\mathrm{p}])$$
$$-ch.\,(([\mathrm{SOX9\ mRNA}].11.921.\,[\mathrm{OtherTargets}])$$
$$-\mathrm{ch}.\,([\mathrm{SOX9mRNA}].59.112.\,[\mathrm{CAV\ 1}])]$$

$$\frac{d[SOX9]}{dt}=[+ch.\,(604.499\ .\,[SOX9\ mRNA])$$
$$-\mathrm{ch}.\,(0.376.\,[SOX9])]$$

$$\frac{d[SOX9PhosphoProtein]}{dt}=[+ch.\,(1.848)$$
$$-\mathrm{ch}.\,(0.00475.\,[\mathrm{TGFB3}])]$$

$$\frac{d[TGFB3]}{dt}=[+ch.\,(6.085.\,[\mathrm{SOX9}])$$
$$-\mathrm{ch}.\,(0.376.\,[0.376.\,[TGFB3]]]$$
